# Statistical Mechanical Prediction of Ligand Perturbation to RNA Secondary Structure and Application to the SAM-I Riboswitch

**DOI:** 10.1101/461749

**Authors:** Osama Alaidi, Fareed Aboul-ela

## Abstract

The realization that non protein-coding RNA (ncRNA) is implicated in an increasing number of cellular processes, many related to human disease, makes it imperative to understand and predict RNA folding. RNA secondary structure prediction is more tractable than tertiary structure or protein structure. Yet insights into RNA structure-function relationships are complicated by coupling between RNA folding and ligand binding. Here, we introduce a simple statistical mechanical formalism to calculate perturbations to equilibrium secondary structure conformational distributions for RNA, in the presence of bound cognate ligands. For the first time, this formalism incorporates a key factor in coupling ligand binding to RNA conformation: the differential affinity of the ligand for a range of RNA-folding intermediates. We apply the approach to the SAM-I riboswitch, for which binding data is available for analogs of intermediate secondary structure conformers. Calculations of equilibrium secondary structure distributions during the transcriptional “decision window” predict subtle shifts due to the ligand, rather than an on/off switch. The results suggest how ligand perturbation can release a kinetic block to the formation of a terminator hairpin in the full-length riboswitch. Such predictions identify aspects of folding that are most affected by ligand binding, and can readily be compared with experiment.

## 1 INTRODUCTION

Ribonucleic acid (RNA) folding is intimately connected with its biological functions. Many of these functions involve interactions with ligands (1). As the known range of biological roles for RNA has expanded-including, for example, enzyme catalysis and diverse mechanisms of regulating gene expression-the ability to predict folding from base sequence has become increasingly important (2).

However, as with proteins, folded RNAs often function in partnership with other cellular components, such as proteins, other nucleic acids, or small molecules. These interactions often perturb RNA folding in a manner that cannot be ignored when attempting to explain biological function. For example, the ribosome undergoes a series of conformational changes that are coupled to a series of binding and dissociation events involving translation factors (3). Thus, the issue of ligand binding is tightly coupled to the dynamic nature of RNA folding. The latter coupling, in turn, is essential for understanding how RNA performs its diverse biological roles. Therefore, it would be desirable to predict not only RNA secondary structure, but also to compute the perturbation to this dynamic folding in response to ligand binding. Programs have been presented to predict the effect of binding by oligonucleotide ligands, with focus on the accessibility of RNA segments, but not, to our knowledge, on binding by non-nucleic acid ligands (4,5).

Even more important is to have a rigorous, quantitative, testable, and preferably simple conceptual framework for understanding how ligand binding is coupled to RNA flexibility. Currently the issue is usually discussed in terms of qualitative descriptive models such as “lock and key”, “induced fit” and “conformational capture” (6–8). These descriptive models and computationally intensive atomistic calculations (9,10), as well as crude virtual screening scoring functions (11,12), are problematic for predicting all but the smallest local conformational changes. The same is true for Molecular Mechanics Poisson-Boltzmann Surface Area (MM-PBSA), Free Energy Perturbation (FEP) and related continuum approximation methods for calculating binding energies (11,13,14). The relative feasibility of calculating plausible free energies for RNA secondary structures, on the other hand, offers an alternative approach to understanding large scale conformational dynamics and relating them to ligand binding.

Ab initio RNA structure calculations are hampered by the fact that complex tertiary interactions influence secondary structure (15–17). In spite of these difficulties, ab initio RNA secondary structure prediction generally produces a more suitable starting point for alignment-based secondary structure prediction than is the case for proteins (18,19). In many cases, reasonable hypotheses for the secondary structure can be generated based on empirical values for modular components of the RNA, such as nearest neighbor sets of base pairs (20). Experimental studies of RNA structures, including chemical reactivity maps, provide rigorous tests of RNA secondary structure predictions (18,21,22). On the theoretical side, standard prediction of the base pairing arrangement of a single sequence identifies the lowest (minimal) Free Energy structure (MFE) and suboptimal structures of slightly higher predicted energy (23). This set can then be readily narrowed based on sequence alignments and co-variation analysis. Thus, if one initiates one’s prediction limited to the secondary structure of the unliganded RNA, the chance of success is strong, especially with sequence alignment data or sparse structural data. This prediction can then provide a good basis for the more challenging task of tertiary structure prediction (16). It also provides a strong starting point for incorporating ligand binding into RNA structure prediction.

Bacterial riboswitches present an ideal model system to extend RNA secondary structure calculations to the prediction of secondary structure of RNA molecules in complex with ligands. Riboswitches have been found most often in bacteria, where they usually lie in the 5’ Untranslated Regions (5’-UTR) upstream of biosynthetic genes (24–26). They contain a bio-sensing element, called the aptamer, which binds a signaling molecule, and a so-called expression platform. The riboswitch undergoes a perturbation in secondary structure upon ligand binding, leading to significant changes in expression of downstream genes (27–30).

Riboswitch thermodynamic and kinetic behavior has been modeled using a systems approach, incorporating coupled rate equations for binding and folding (31–35). These simulations assumed a binding constant for the ligand to the aptamer-forming structure, which could be estimated based upon biophysical measurements. This type of formalism also allows for a (presumably reduced) binding affinity for the alternative conformer(s), which are presumed to predominate in the absence of ligand. To our knowledge, no previous small molecule binding prediction took account of differential binding affinities to the populations of intermediate conformers (36). Yet such differential affinity could significantly affect the population distribution of conformers at equilibrium.

Using the well-studied S-Adenosyl Methionine (SAM)-I riboswitch as a model system, we hypothesized that a range of secondary structures could play a role in ligand-induced refolding (30,37). Riboswitch fragments designed to mimic these intermediate structures were shown to bind to SAM but with reduced affinity compared to the fragments which have been truncated to form the aptamer fold exclusively (38). These findings gave rise to potential thermodynamic and kinetic mechanisms for coupling SAM binding to riboswitch secondary structure (30).

In this study, we describe a statistical mechanical formalism to simulate and predict the impact of a small molecule ligand on RNA population. As far as we know, no previous study has used a rigorous statistical mechanics approach to thermodynamically simulate ligand induced conformational changes in RNA. We apply the method on the SAM-I riboswitch incorporating previous experimental findings to secondary structure calculations for the *B. subtilis yitJ* SAM-I riboswitch in the presence and absence of SAM. As in previous work (37), we compute thermodynamic properties at a series of transcript lengths, to simulate co-transcriptional folding. In this work, rather than calculating base pairing probabilities, we directly compute the probabilities of secondary structure folds at the range of transcript lengths, for which thermodynamic calculations suggest a “sensing window” (37,39), in the presence and absence of SAM. The aim of this study is to determine the effect of the ligand (SAM), on the SAM-I riboswitch conformational population distribution. Here, we focus on how the ligand influences the population, including strand migration intemediates, at increasing RNA lengths within the decision window.

The results indicate a significant increase in the fraction of conformations containing a nucleated P1 helix, an important component of the aptamer secondary structure, in the presence of SAM, but only for a narrow range of transcript lengths. The study illustrates that simple statistical thermodynamic models can be used to determine the effect of a ligand on the energetics of a RNA population, and quantitatively simulate the conformational shifts associated with the ligand binding.

## 2 METHODS

In this work, we introduce a formalism describing the effect of the presence and absence of the ligand on the RNA population, as a thermodynamic product and without considering kinetic barriers.

### 2.1 The energetic description of the system

A macromolecule-ligand interaction, such as the interaction between RNA and a ligand, can be represented as:

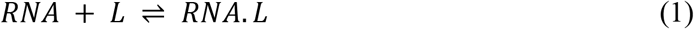

*where*, *RNA*, *L* and *RNA*. *L* are the concentrations of the RNA, the ligand and the RNA-ligand complex, respectively. At equilibrium the reaction free energy ∆*G*, is 0, and we can write:

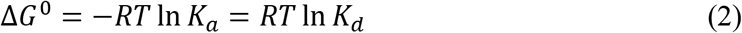

*where*, ∆*G*^0^ is the molar free energy difference at standard conditions i.e. at *T* = 298.15 Kelvin and at 1 atm pressure. The free energy of the system is described relative to a reference state (an RNA in the same conformation plus the free ligand). *R* is the gas constant, and the equilibrium association constant, *K_a_*, represents the equilibrium ratio between the concentrations of the products and the reactants, which is given by,

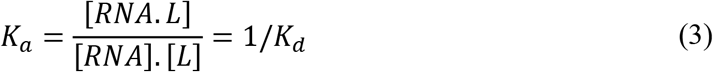

*where*, *k_a_* and *k_d_* are the association and dissociation constants, respectively, and the [ ] notation denotes the concentrations. Upon binding with ligand, the equilibrium free energy difference for the system of RNA plus ligand, relative to the reference state of free RNA of the same conformation plus free ligand, can be described by:

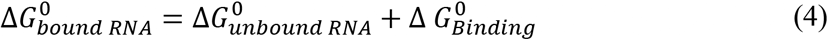

*where*, 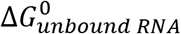 and 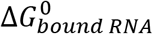 are the free energy of the RNA population *before* and *after* binding to the ligand, respectively. The quantity 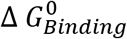 is the total free energy change associated with ligand binding. In fact, the last term in equation 4 refers to a free energy change for a bimolecular system, and thus includes a contribution from the change in free energy of the ligand upon binding. The free energy difference is independent of the ligand concentration and RNA conformer population.

The above treatment assumes that the RNA forms two conformations, or that any variation among bound comformations does not affect ligand affinity. To account for differential binding to RNA conformers, a statistical mechanical treatment, incorporating a partition function, is required. The treatment above allows us to compute altered folding free energies for the individual conformers in the presence of the ligand. The altered free energies for individual conformers are incorporated into the statistical mechanical treatment described below.

### 2.2 Determining the RNA population: probabilities of the RNA conformers in the presence and absence of the ligand

The partition function (40,41) *Q* for *N* conformers, assuming neglegible ligand-ligand interaction, is generally considered to be decribed by the equation,

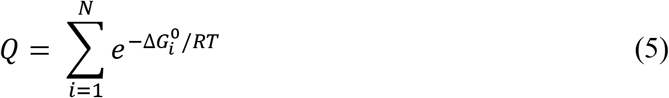

*where*,

Q is the partition function,
∆*G_i_* is the free energy of the conformer *i* in the units of *kcal/mol*,
*R* is the Gas constant
*T* is the temperature in Kelvin

Then, *P_i_* which is the probability of conformer *i* is given by,

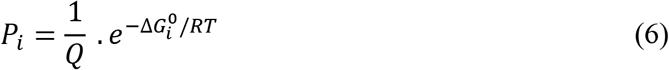

In the absence of the ligand (*before* binding), the partition fuction *Q_BB_* is thefore given by,

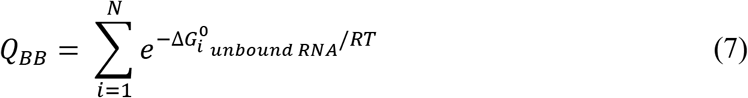

*where*,

*Q_BB_* is the partition function for the system *before* binding,
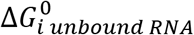 is the folding free energy of the conformer *i before* binding, in the units of *kcal/mol*, as calculated by the *RNAsubopt* program.

Then, *P_i BB_* the probability of conformer *i before* binding is given by,

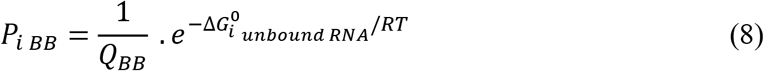

To determine the probabilities of the RNA population in the presence of the ligand (*after* binding), the partition function is re-calculated using the modified free energies after the binding from the equation,

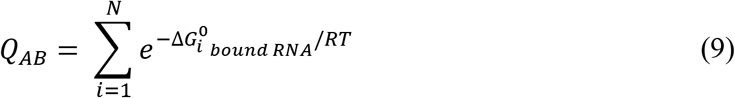

*where*,

*Q_AB_* is the partition function for the system *after* binding,
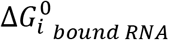 is the free energy difference of the conformer *i after* binding, in the units of *kcal/mol*.

Then, the probability *P_i AB_* of conformer *i after* binding is calculated from,

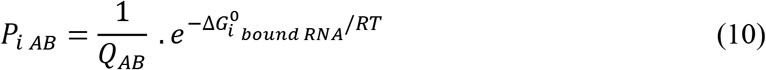

The modified/ re-scaled free energies 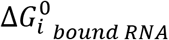 are obtained according equation (4).

It should be noted that, since the focus of the calculation is purely on the RNA conformer populations in the bound state, the relative values of 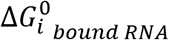 will determine the outcome. The value of 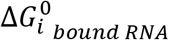 is independent of the total concentration of reactants. Thus, the calculation does not account for the overall ligand concentration dependence for the conformer population distribution. It does, however, provide a rigorous description of the conformer distribution for the bound subpopulation of RNA (the population “after binding”). The quantity 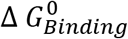 is assumed to be equal to the summation of free energy contributions of both specific and non-specific binding.

If ligand binding is associated with a specific substructure, then the set of conformers containing this substructure can be considered as a macrostate. The probability of this macrostate *before* binding to the ligand is calculated as the summation of all probabilities of the constituting microstates,

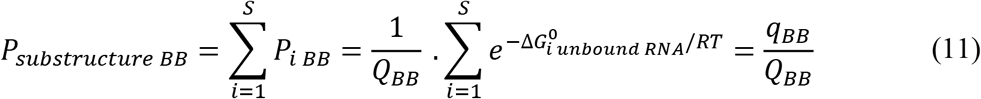

*where*,

*Q_BB_* is the partition function *before* binding with the ligand.
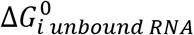 is the free energy of conformer *i before* binding with the ligand.

q_*BB*_ is the sum of the Boltzmann factors for conformers that contain a particular substructure *before* binding with the ligand. The summation in the equation above is over *S* states, which contain that particular substructure. The probability of this macrostate *after* binding is calculated from,

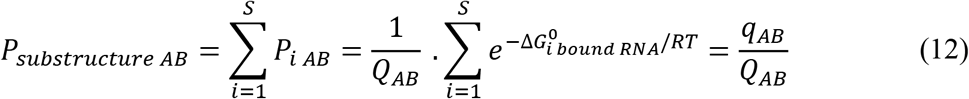

*where*,

*Q_AB_* is the partition function *after* binding with the ligand.
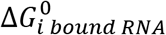 is the free energy of conformer *i after* binding with the ligand.
*q_AB_* is the sum of the Boltzmann factors for conformers that contain a particular substructure *after* binding with the ligand.

### 2.3 Constructing free energy landscapes of macrostates in the absence and presence of the ligand

A free energy landscape is constructed by calculating the macrostate free energies from groups of conformers having a specific substructure in the presence and absence of the ligand. The free energy landscape for macrostates was constructed from their corresponding probabilities (obtained from equations 11 and 12, respectively). The corresponding free energy of a substructure or macrostate in the absence and presence of the ligand is calculated from,

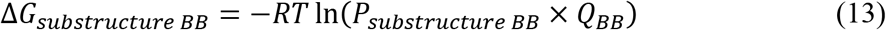

and,

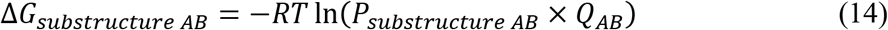

### 2.4 Application to the SAM-I riboswitch

#### 2.4.1 Calculating the binding energies of SAM to the SAM-I riboswitch

The method described above was applied to study the effect of SAM on the *yitJ* SAM-I riboswitch. As described in the Supplementary Methods, the interaction of the RNA with SAM can be represented by the thermodynamic cycle involving RNA in structured or unstructured states, and in forms where it is bound or unbound to SAM (**Figure S1**).To test the robustness of the results with respect to uncertainties in experimental equilibrium dissociation constants (see Supplementary Note 1) and to evaluate the impact of the differential binding between the aptamer and intermediates, calculations were repeated to obtain probability distributions using different sets of binding energies, 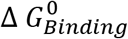. The binding energies were obtained from single titration point equilibrium dialysis data, from Scatchard plots derived from multiple equilibrium dialysis measurements (38) or from two sets of data for hypothetical ligands of stronger binding affinity (hypothetical ligand 100x and hypothetical ligand 1000x).

For single titration point data, the probabilities presented were obtained using an energy range of 8 kcal/mol from that of the MFE from data in (38) as described in the Supplementary Methods. For the Scatchard plots and hypothetical ligands, all intermediates were assumed to bind with similar affinities to the ligand and the probabilities were also obtained using the same energy range of 8 kcal/mol from optimum. Specifically, in the case of Scatchard plots, the *K_d_*s for all intermediates were assumed to be equal to that which was measured for *3P1_10AT* (38), which is the only intermediate for which binding was measured using this method. In all cases mentioned above, 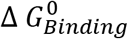 was assumed to be equal to:

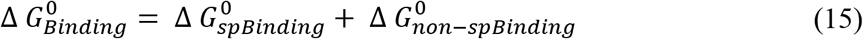

where 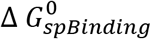 and 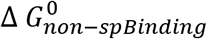 in equation 15 are the SAM binding free energies to the riboswitch in the presence and absence of the P1 helix structure, respectively. Previous studies suggested that, the presence and stability of the P1 helix had been correlated with SAM binding (38, 42–44). Thus, in our calculations, we assumed the free energy of binding of a construct that lacks the P1 helix (*P2-AT* in reference (38)) to represent the non-specifc binding 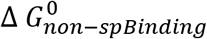. Since all RNA may have non-specific binding, it has no effect on the calculated probabilities after binding, and it was assumed to be the same across all binding data sets. Hence, only the specific binding 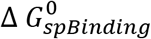 was used in the energy re-scaling i.e. in obtaining 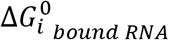

#### 2.4.2 Calculating the probabilities of intermediates and transition states in the presence and absence of SAM

The predicted RNA optimal and suboptimal secondary structures were scanned for the presence of substructures that are hypothesized, based on data with analogue truncated molecules (38), to be capable of binding to SAM. These structures therefore are assumed to manifest a modified free energy in the presence of the ligand. Each conformer was classified to belong to either one of the following:

1. Strand Migration Intermediates (binds to SAM), illustrated in **Figure 1**: *2P1_11AT, 3P1_10AT, 4P1_9AT, 5P1_8AT, 6P1_7AT, 7P1_6AT, 8P1_5AT* Here numerals refer to the number of base pairs in the competing P1 and “anti-terminator” (AT) helices, respectively. The antiterminator helix takes its name due to competition with the downstream rho-independent terminator helix, in the absence of ligand (45–47). The P1 helix is a critical component of the four-helix containing SAM-binding aptamer, and potentially blocks the formation of the AT helix, facilitating termination of transcription. Two sets of calculations were made using a search criteria *“continuous AT”* and *“discontinuous AT*”. In the latter, one mismatch/unpaired base was allowed in the AT stem. For example *2P1_11AT* and *2P1_10AT* were considered indistiguishable. Hence, *2P1_10AT, 3P1_9AT, 4P1_8AT, 5P1_7AT, 6P1_6AT, 7P1_5AT, 8P1_4AT* were assumed to bind to SAM with similar binding energies to the corresponding intermediate with continuous/fully base paired AT loop. Here *XP1_YAT* includes any substructure with *Y AT* base pairs, regardless of their distribution. In other words, the single unpaired base may occur anywhere within the AT stem. Additional structures within this category include those with base pairs within the AT loop (**Figure S2**, panels **C** and **F**), which are assumed to be present within the experimental binding data, and therefore are included with the same binding free energy.
2. Transition States *Aptamer_TS* (binds to SAM), illustrated in **Figure 1**, which contains a full P1 helix but only a partially base paired AT loop (less than 4 base pairs). In the absence of the ligand, this group of intermediates are less stable compared to those containing *8P1_5AT* (see below). Based on analogy to binding data for the truncated aptamer (38,46), however, this group is assumed to bind SAM with more favourable free energy.
3. Any conformer that does not belong to one of the two groups above is considered to not bind to SAM.

**Figure 1.**
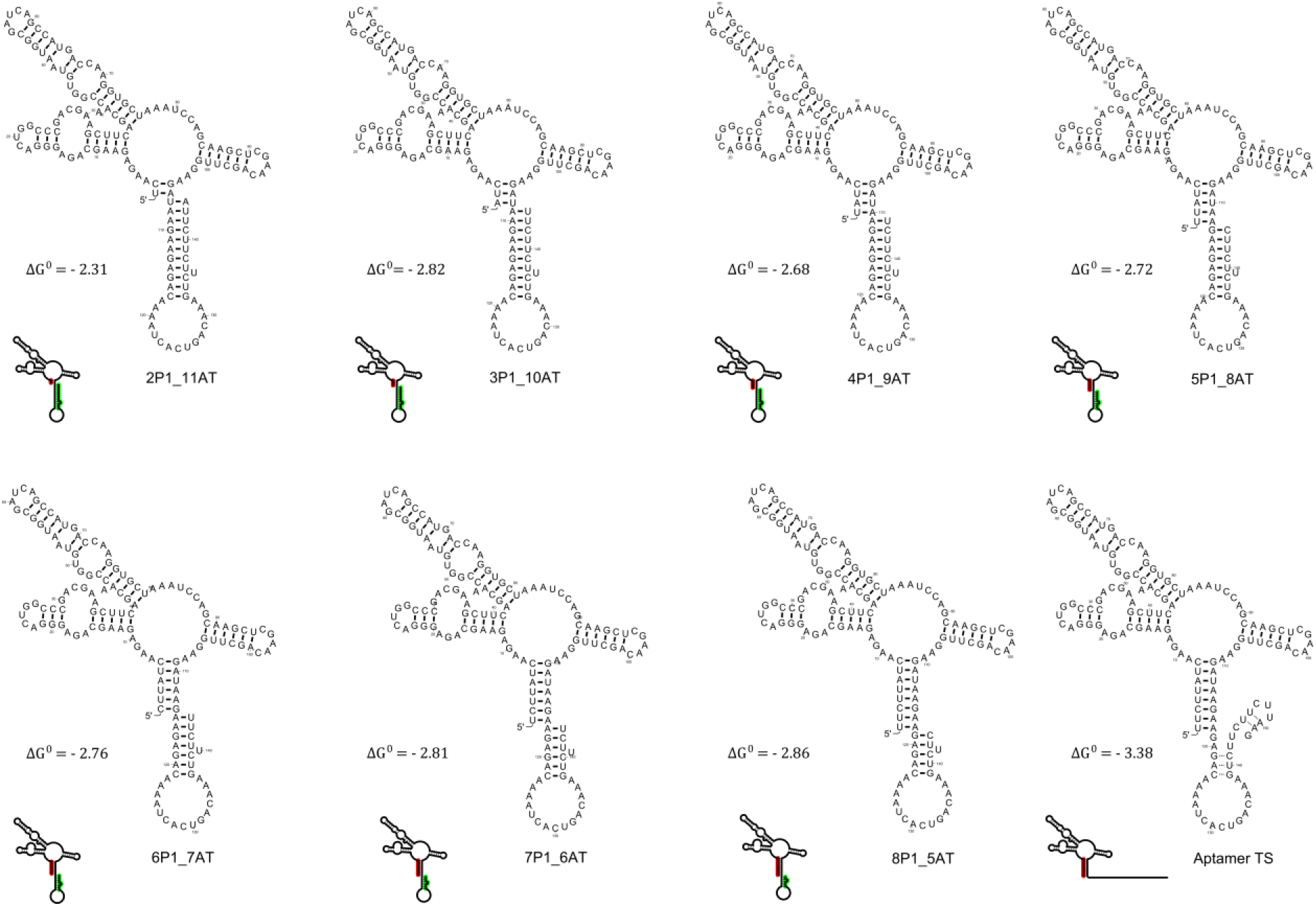
Secondary structures and skeleton representations of the proposed strand migration intermediates. The figure illustrates examples of secondary structures and skeleton representations of constructs/truncated intermediates as well as the aptamer transition state (*Aptamer_TS*) along with their specific binding energies (∆*G*^0^) at standard conditions, in the units of kcal/mol. The dotted lines in the *Aptamer_TS* represent examples of possible base pairing. The convention for naming the conformers is XP1_YAT, where X is the number of base pairs formed in the P1 helix, an important component of the SAM-binding aptamer (containing a bulged four-way junction). Y is the number of anti-terminator (“AT”) base pairs formed. The AT helix is considered a competitor with the P1 helix. Thus, where X≠ 0, Y≠ 0, we have a binding-competent strand migration intermediate.

The first two categories are classified as “binding-competent” conformers (48), while the third category is considered to be “binding-incompetent”. RNA structures which contain a particular substructure (that is, a specific configuration within the the P1/AT competition region) were considered indistinguishable, regardless of the configuration in other parts of the molecule (i. e. helices P2-P4 and linking segments), and likewise for those containing the *Aptamer_TS*. Such “indistinguishable structures” (indistinguishable from the available binding data) were lumped into a single macrostate. The probability of this macrostate was calculated as the summation of all probabilties of the constituting states according to equation 11. In the case of the SAM-I riboswitch, for example, the substructure may exist alongside variations in secondary structures in the P4 and in parts of the P2 and P3 helix regions, with limited effect on measurable binding parameters.

A string search through the suboptimal RNA conformers using both their secondary structure bracket notation and the corresponding sequence/position was used to identify the occurrence of each substructure. In order to account for strand “slippage” and base pairing variations in the AT loop (**Figure S2**), counting the number of base pairs in the AT loop was made flexible, in such a way that it allowed base pairing anywhere in the AT loop. As described earlier, the corresponding energies of conformers that bind with SAM were re-scaled, while those that do not bind with SAM remained unchanged. The resulting new set of energies 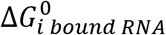 were used to calculate the probabilities of substructures after binding from equation 12. The corresponding free energies of macrostates determined by groups of conformers having different number of (competing) base pairs in the P1 and AT loops, in the absence and presence of SAM, were calculated from equations 13 and 14, respectively.

### 2.5 Implementation of the method using *RNAsubopt*

The optimal (mimimal free energy or MFE) structure and the suboptimal structures (higher energy structures compared to the optimal within a specified energy range) of the SAM-I riboswitch were calculated, for a range of increasing transcript lengths. Optimal and suboptimal RNA secondary structures were calculated using the Vienna package *RNAsubop*t software (49). For all calculations throughout the manuscript, *RNAsubopt* options were set to *–e* 8, i.e. suboptimal structures within an energy range of 8 kcal/mol from the MFE unless otherwise stated, and *-T* to 25°C i.e to room temperature 298 Kelvin. Some simplifying assumptions were incorporated in order to focus the analysis on the effect of ligand binding on the RNA conformational distribution. Specifically, to reduce the number of conformers, the option *– noLP* was used, implying that no helices with only a single base pair were allowed. Default parameters were used otherwise. It should be noted that the default parameters in the *RNAsubopt* program apply a simple model for the treatment of the dangling ends and usually ignore coaxial stacking. Namely, in the default option for the dangling ends and coaxial stacking treatment (option *-d2*), the energetic contributions of the stacking of an unpaired dangling base, present between two helices, are added on both sides of the base, whereas coaxial stacking of helices are ignored. A more sophisticated treatment of coaxial stacking needs be investigated in the future (e.g. (50) and references therein). RNA structures were plotted using the *RNAplot* program (51).

## 3 RESULTS

We wished to derive a method for simulating conformational shift/rearrangement in the RNA population, incorporating the effect of bound ligand. The method was applied to study the effect of SAM on the *yitJ* SAM-I riboswitch. Starting with calculated free energies for suboptimal secondary structures, we searched those secondary structures for motifs that have been shown to correlate with ligand binding. The method then modifies the free energies of those binding-competent suboptimal structures to obtain a new set of energies for the RNA population. Through the calculated partition function before and after binding, the probabilities of conformers containing a specific secondary substructure or motif can be estimated in the presence and the absence of the ligand. The method bridges the gap between conformational capture and induced fit by making rigorous quantitative prediction of individual conformer populations probabilities.

### 3.1 The ranges of dissociation constants and binding energies of the SAM-I riboswitch aptamer and proposed intermediates

The binding constants for “strand migration intermediates” (see Materials and Methods section, **Figure 1** and **Table S1** for definitions of those intermediates) were determined from single titration point equilibrium dialysis measurements (38) and calculated as described in the Supplementary Methods. The *K_d_* values for constructs representing intermediates (**Figure 2** and **Table S2A)** ranged from ~ 0.79 − 2.0 micromolar (*μ*M), for the tightest (*8P1_5AT*) and weakest (*2P1_11AT*) binding intermediates, respectively. The latter *K_d_* values correspond to total binding free energy differences at standard conditions 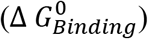 ranging between −7.77 to −8.32, kcal/mol. Constructs *P2_AT* and *Aptamer* (*“Aptamer_Mg_snap_cooled”*) had *K_d_* values of 9.99 × 10^−5^ M and 3.30 × 10^−7^ M, respectively. Thus, the *Aptamer* construct binds ~ 300-fold more tightly than the *P2_AT*. The latter *K_d_* values correspond to standard binding free energies 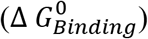 of −5.45 and −8.84 kcal/mol, respectively. The difference of ~ −3.38 kcal/mol therefore represents the binding energy in the presence of a fully base paired P1 and in the absence of a competing strand from the AT loop. A more comprehensive list of binding energies for other constructs are compiled in **Table S3**.

**Figure 2.**
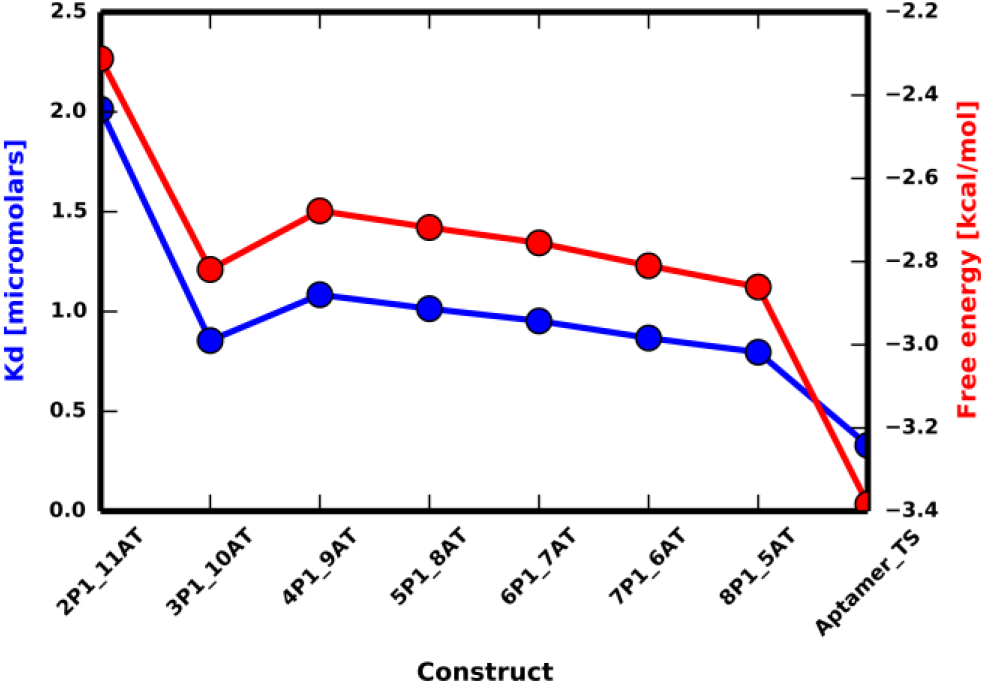
Dissociation constants and specific binding free energies of the truncated strand migration intermediates. The figure shows a plot of dissociation constants (*K_d_*), depicted in blue, and their corresponding *specific* binding free energies, depicted in red, for constructs used in this study. The constructs were designed to represent strand migration intermediates. The dissociation constants shown here were calculated from single titration point equilibrium dialysis data, measured by Boyapati et al (38). The free energies were calculated from the dissociation constants and after subtracting the *non-specific* binding energies from the total binding energies. The resulting *specific* binding free energies can therefore be correlated with the presence and stability of the P1 helix. The uncertainty in 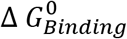 (due to reported random error in the primary data) ranges from 0.011 to 0.064 kcal/mol, except for *3P1_10AT* for which the uncertainty in 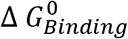 is equal to 0.24 kcal/mol, (see Supplementary Note 1 for a discussion on the uncertainties in those measurements).

Upon the subtraction of the *non-specific* binding energy 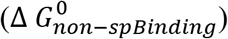 to obtain the P1 *specific* binding energy contribution 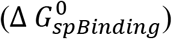 for each construct (which could be defined as the energetic cost upon binding due to the presence of the P1 helix for each construct), energies with magnitudes ranging between −2.31 and − 2.86 kcal/mol for the strand migration intermediates and −3.38 kcal/mol for the *Aptamer* (**Figure 2**) were obtained. This variation in binding free energy among the binding-competent intermediates is of the order of ~ 0.55 kcal/mol, while the difference between the intermediates and the *Aptamer* ranges from ~ 0.52 to ~1.07 kcal/mol. This free energy difference trends towards stabilization of the conformers containing the fully folded P1 helix (and in the absence of a competing strand from the AT loop) relative to those with only a partially base paired P1 and/or base pairing competition exerted by a strand from the AT loop. Thus, if this equilibrium were reached during the respective window during transcription, it would tend to prime the system for the downstream unfolding of the AT loop.

Scatchard plots, that are based on equilibrium dialysis measurement, can provide more accurate *K_d_* measurements, since the values are deduced from numerous measurements as opposed to single titration point measurements. A small margin of difference had been reported in the deduced *K_d_* from the two techniques (38). To test if this difference would affect the results, we performed the calculations using both datasets independently. The limitation of the Scatchard plots binding data is that data points for only two constructs, namely the *3P1_10AT* and the aptamer were available in the previous study (38). The *K_d_* values derived from Scatchard plots and used for probability calculation were 7.90 × 10^−7^, 9.99 × 10^−5^ and 3.20 × 10^−8^ M for the intermediates, *P2_AT* and aptamer, respectively (**Table S2B**). The latter values reflect a superior discriminatory power (higher differential binding) in the favour of the aptamer over the intermediates, if compared with single titration point data. The corresponding specific binding energies 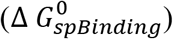 for the intermediates and aptamer were −2.87 kcal/mol and −4.76 kcal/mol, respectively. Calculations were also performed using two hypothetical ligands (hypothetical ligand 100x and 1000x). The *K_d_* values were set in such a way that they would exhibit an even higher differential binding, to determine what differential binding would result in the dominance of the *Aptamer_TS* structure. The *K_d_* values for these hypothetical ligands and their corresponding binding energies are tabulated in **Table S2C** and **Table S2D**, respectively. A tenfold decrease in *K_d_* is predicted to tip the balance towards the *Aptamer_TS* structure but intermediate structures become negligible only with another order of magnitude in differential affinity.

### 3.2 Broader probability distributions in the absence of SAM are observed at longer riboswitch lengths (≥ 150 nucleotides)

Computing the populations of all possible secondary structures of a riboswitch of well over 100 nucleotides is computationally expensive. On the other hand, conformations with free energies above a certain threshold will make a negligible contribution to the overall ensemble. We chose to include conformations with free energies up to 8 kcal/mole higher than that of the MFE, corresponding to probabilities of suboptimal conformers that are approximately six orders of magnitude lower than that of the MFE in the temperature range 298-310 K. The number of suboptimal structures within the tested energy range increases as a function of length and ranges from ~3.73 × 10^4^- ~2.48 × 10^5^ structures for lengths 145 - 152 (**Figure S3A, Table S4**). From the perspective of this study, we are not concerned of the details of these structures, but we need to distinguish between binding-competent and binding-incompetent structures. At a transcript length of 145 nucleotides, the number of conformers found within the 8 Kcal/mol range relative to the MFE and that are binding-competent is approximately ~ 2.82 × 10^4^conformers, and this number increases to reach ~ 5.37 × 10^4^ conformers at sequence length 149 nucleotides (highest), and then the number of binding-competent conformers sharply decreases again at 150 or longer (**Figure S3B**). Thus, the number of different conformers that are capable of binding to SAM within the selected free energy window of the MFE peaks at 149. This number of conformers alone is not a measure of the amount of SAM bound. First, the number of possible suboptimal structures increases with longer transcription/RNA lengths and second, the stability of each conformer must be weighted according to the corresponding Boltzmann factor, based on the individual conformer free energies. **Figure S3C** shows the binding-competent conformers as a fraction of suboptimal structures within the free energy window, as a function of transcript length. The largest fraction of conformers within the free energy range that are binding-competent (~ 76%) was at length 146 nucleotides from the beginning of the riboswitch.

Histograms of the energy distribution of the RNA population (**Figure S4**) show a smooth distribution of free energies of the population at all examined transcription lengths in the absence of SAM. In order to predict the binding-competent proportion of the population at a given time, we went on to calculate the total probability of conformers having one of the substructures, at a given transcription length. The explicit description of the conformer population is a unique feature of the current approach, which provides crucial insight into the perturbation of the Free Energy Landscape due to ligand binding. In the absence of SAM, the sum of probabilities of all binding-competent conformers decreased from 98% to 7% at transcription lengths from 145 to 152 (**Figure 3**, **Table S5**). The binding-competent strand migration intermediates, dominate at lengths 145-147, and still represent a significant proportion of the population at transcript length 148. A bigger decrease is seen at 149: only 35% of the population can bind to SAM. An abrupt reduction, however, of the total probability of the population that can bind to SAM was observed at a transcript length of 150 (or longer). Further, at this length (≥ 150 nucleotides) a broader probability distribution is observed compared to shorter lengths (**Figure S5**). This result agrees with previously reported calculations (37) that the unbound riboswitch favours AT helix formation in a short transcription window starting near a length of 150 nucleotides.

**Figure 3.**
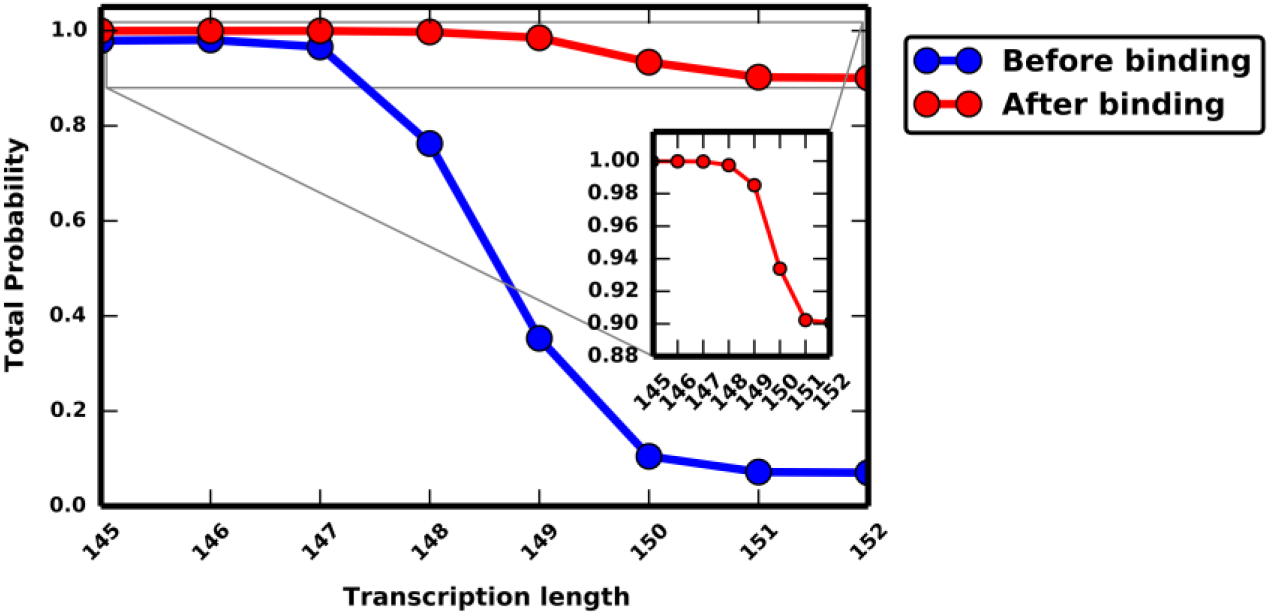
The predicted cumulative probability of all binding-competent conformers *before* and *after* binding with SAM at various transcription lengths. The plots show the summed probabilities of all binding-competent conformers *before* binding with SAM (i.e. in the absence of SAM, blue) and *after* binding with SAM (i.e. in the presence of SAM, red). The inset plot of the total probability of binding-competent conformers *after* binding illustrates that a decline in the total probability is predicted at longer transcription lengths. The total probabilities were obtained by summing the probabilities of intermediates and *Aptamer_TS* in each of the two conditions at a given transcription length.

The number of different conformers per intermediate, and corresponding fractions are tabulated in **Table S6** and illustrated in **Figure S6**. In the absence of SAM, the riboswitch population is dominated by a fully base paired P1 helix (in the unbound state) up to transcript length 148. The probability of conformers containing the *8P1_5AT* substructure, i.e. with a fully base paired P1 helix, represented ~ 83.8% at transcription length 145 and gradually declined with the increase of transcription length to reach ~ 4.56% of the population at transcription length of 152 nucleotides (**Table S5**, **Figure S7)**.

### 3.3 The effect of SAM on the RNA population: The presence of SAM stabilizes the intermediates with fully folded P1 helix

Calculations were made using binding energies obtained from two experimental and two simulated (hypothetical) sets of binding data. The reader may refer to the Materials and Methods section for a detailed description of the energetics of the system and the probability calculations. **Tables S2** (**A**, **B**, **C** and **D**) show the binding data used and the corresponding free energy of binding. These values were added to calculated folding free energies to obtain the probabilities after binding (i.e. probabilities in the presence of the ligand).

We start by presenting results incorporating binding free energies derived from single titration point data. Histograms of the energy distribution of the RNA population (**Figure S8**) show that the presence of SAM in the environment sharply perturbs the free energy distribution (or free energy landscape) of the SAM-I riboswitch population, by lowering the free energy of only a subset of the population (compare to **Figure S4**). This selection process is reminiscent of the so-called conformational capture mechanism, in the sense that pre-existing conformers are stabilized upon binding, but differs in that multiple conformers are still present (30). The gap observed between two clusters in the histograms becomes less abrupt at transcription length 150 or longer. This gap represents the difference in free energy between binding-competent and binding-incompetent conformers.

**Figure S5** shows plots of energies versus probabilities of the SAM-I riboswitch, at lengths 145-152, before and after binding with SAM, hence illustrating the impact of SAM on the population. The analysis of the probability distribution after binding showed that at transcription lengths 145, 146, 147, 148 and 149 the total probability of binding-competent conformers (or bound conformers i.e. containing one of the substructures/intermediates) range between ~0.90 and ~1.0 (**Figure 3** and **Table S5**) when SAM is assumed to be bound. A modest but significant reduction (6-10%) of the total probability of bound conformers at transcription lengths 150 or longer was predicted. Specifically, at transcript lengths of 150, 151 and 152, the total probability of conformers containing the intermediates/substructures were 0.93, 0.90 and 0.90, respectively.

Conformers having the *8P1_5AT* substructure have the lowest energy and hence constitute the majority of the population after binding with SAM. Hence, the presence of SAM can be correlated with the presence of the fully base-paired P1 helix. Specifically, the calculations showed that, in the presence of SAM, the percentage of conformers containing the *8P1_5AT* substructure declines from 84.6% to 61.1%, for transcription lengths 145-152 nucleotides (**Table S5**, **Figure S9**). A consequence of the gradual reduction of the percentage of conformers with fully base paired P1 helix is that the downstream formation the aptamer with a termination loop, which is responsible for transcription termination, will require unpairing of the AT helix. While a sharp decline is particularly seen at transcription length 150 or longer in the absence of SAM, the decline is much less in the presence of SAM. It is clear, that the effect of SAM on the population before transcription in the length range of 148 to 150 is more dramatic compared to the limited effect after this transcription length (**Figure 3** and **Figure S5**).

Comparing the probabilities of conformers in the tables *before* and *after* binding to SAM shows that the binding of SAM leads to 1) increase of the population of conformers that have a nucleated P1 helix, particularly in the “decision window” between 150-152 nucleotides, 2) within the latter set of conformers, an increase in the contribution of those with a full P1 helix to the point that they dominate the population, 3) an increase in the probability of conformers with an unfolded AT loop (*Aptamer_TS*) and 4) conformers that lack the P1 helix (which are binding-incompetent) are reduced to a minority of the population. This stabilization of the P1 and unfolding of the AT loop subsequently will lead to lowering of the activation energy required for the formation of the termination loop once its sequence is transcribed downstream. It should be noted that, at any given length in the studied window, if only AT base pairs as defined in the expected full length AT helix (i.e. as shown in **Figure 1**) are included in the binding-competent category, and if a *continuous AT* loop is required, then the total probability of the substructures before binding, even at lengths 148 or 149, is found to be low (data not shown). Since still a small percent of the RNA population incorporates mismatches and “slippage” in the AT loop, and these are not likely to affect binding, the *discontinuous AT* set of probability calculations is likely to be more accurate. As reported in the Materials and Methods section, this assumption has been incorporated into the results reported here. The effect of this assumption is described in detail in section 3.6.

Similar trends were observed upon using binding energies obtained from Scatchard plots (**Figure S10** and **Table S5**). The latter calculations yield up to ~ 2.3% increase in the total probabilities of the binding-competent conformers after binding (last column in **Table S5**). This difference comes from a significant redistribution in the population. Specifically, the *Aptamer_TS* probability significantly increases (up to ~ 23.65%) compared to the calculations using the single titration point data, whereas *8P1_5AT* is reduced by ≤ 21.45%. **Figure 4** illustrates the impact of SAM on the probabilities of the intermediates, including the aforementioned increase of *Aptamer_TS*, at a chosen transcription length 149. This shift reflects a greater differential binding in favour of the aptamer compared to the intermediates in the Scatchard data.

**Figure 4.**
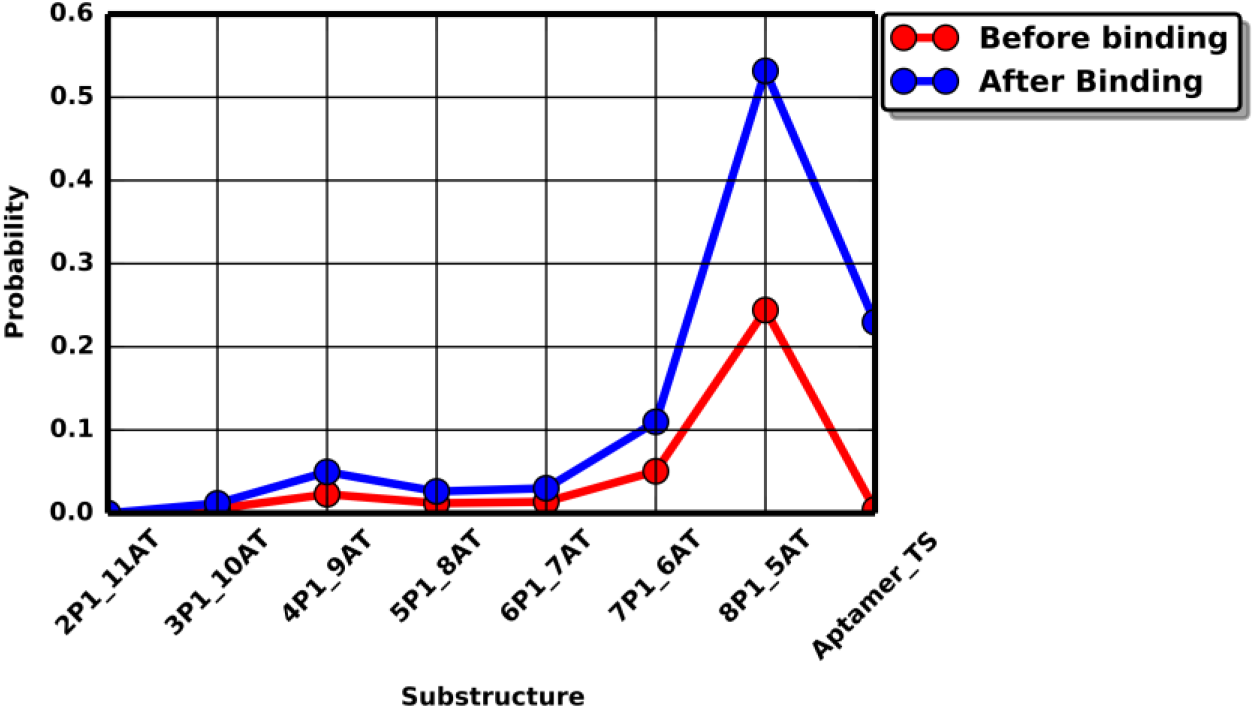
The predicted probability of the strand migration intermediates and the *Aptamer_TS before* and *after* binding with SAM, at transcription lengths of 149, and using the Scatchard plot binding data. The plot shows the probability of strand migration intermediates as well as the *Aptamer_TS* calculated in the *presence* (blue) and *absence* (red) of SAM, calculated using the Scatchard plot binding data, at transcription length 149 bases. The plot illustrates a significant increase in the *8P1_5AT* and *Aptamer_TS* after binding. Both substructures contain a fully base paired P1 helix. The *Aptamer_TS* also contains a fully or partially unfolded AT loop, paving the way for the formation of the termination loop through an accelerated strand migration process. Similar plots for other transcript lengths are shown in **Figure S10**.

Because the *Aptamer_TS* represents conformers that possess an unfolded AT loop, which are therefore primed for formation of the terminator loop, we wanted to determine what extent of differential binding would be required for the *Aptamer_TS to* dominate the population. For this purpose, two hypothetical ligands with a higher differential binding were tested; hypothetical ligand 100x (**Figure S11** & **Table S7**) and hypothetical ligand 1000x (**Figure S12** & **Table S7**). The probabilities after binding showed that the *Aptamer_TS* exceeded other substructures (beyond 50%) in the population with the hypothetical ligand 100x, and before transcription length of 150. The latter, i.e. the *Aptamer_TS*, overwhelmingly dominates the population e.g. > 90% when the ratio between *K_d_* of the intermediates/Aptamer is equal to 10^3^. Altogether, these results indicate that a bigger differential in aptamer binding to SAM than has been observed would be required to complete unfolding of the AT loop before the terminator loop is fully transcribed.

### 3.4 The predicted effect of key mutations on ligand-induced riboswitch folding

The presented simulation method can be used to predict the effect of mutations on the population in the presence and absence of the ligand. **Figure S13** shows the total probability of the binding-competent conformers of the SAM-I riboswitch in the absence and the presence of SAM (panels **A** and **B**, respectively) of various mutants. Mutants that perturb or destabilize the P1 helix are predicted to reduce the probability of binding-competent conformers, both in the presence and absence of SAM. This outcome should inhibit transcription termination. Some mutants increase the stability of the P1 helix compared to the wild type, leading to the predicted increase in the population of binding-competent conformers compared to wild type at all transcript lengths, including lengths of 150 nucleotides or longer. A predicted consequence would be transcription termination even at low SAM concentrations, since the transcript is primed to form the terminator even when SAM is not present.

Interestingly, similar findings have been observed experimentally (46). Mutants that perturbed the P1 helix have been shown to reduce the binding and reduce the termination in both the presence and absence of SAM (46,52). Whereas, mutants that stabilize the P1 helix increased the fraction terminated in both the presence and even in the absence of SAM (52).

### 3.5 Validating the appropriate energy range, sufficient conformation sampling, and appropriate transcription window

To verify that the suboptimal structures of free energies more than 8 kcal/mole higher than that of the MFE make a negligible contribution to the overall results and conclusion, the total probability of all conformers that are assumed to bind to SAM (strand migration intermediates) in the presence and absence of SAM, were calculated at different energy ranges (from 2 - 11 kcal/mol). For this purpose, we calculated the total probability at length 148, since at this length a sufficient proportion of those binding-competent conformers are populated (**Figure S3**). Previous studies indicated that this length allows sampling of conformers representing strand migration intermediates (37). The total probability before and after binding with SAM, both including binding to substructures with an interruption in AT loop base pairing (**Figures S14A** and **S14B,** respectively) and including only those with a *continuously* base paired AT loop (**Figures S14C** and **S14D,** respectively) are presented. The probabilities, before or after binding, showed no significant changes (<< 0.01%), when the energy range was changed from 8 kcal/mol to 11 kcal/mol. This energy range or wider is therefore generally sufficient to estimate the perturbed probabilities to a first approximation. The number of conformers must increase as a function of the energy range and sequence length. Hence, the energy range of 8 kcal/mol can be thought of as a reasonable balance between accuracy and computational time. The latter energy range was used for all calculations, unless otherwise explicitly stated.

Riboswitch structures are normally depicted beginning from the 5’ end of the P1 helix. In this case, the length of the full *yitJ* SAM-I riboswitch is 157 nucleotides. Competition between the AT helix and the P1 starts at transcription length 145. The termination loop can be nucleated, in principle, using a sequence that is 149 bases long (one base pair can possibly form at this length). The window 145 - 152 bases was therefore studied, to determine the effect of SAM on the competition between P1 and AT helix formation. Previous studies had indicated that this length represents a “decision window” for riboswitch folding (37,39). At the same time, the formation of AT base pairs in this window has a downstream impact on the induction/nucleation and fomation of the termination loop.

### 3.6 Probabilities of the *discontinuous* AT loop represent the RNA population more efficiently compared to the probabilities of the *continuous* AT loop

As described in the methods section, the probability calculations for conformers containing the substructures that represent strand migration intermediates were made assuming two different criteria for substructures. The first set of calculations considered a conformer to have a particular substructure if the number of base pairs in both the P1 helix and the AT loop exactly match the specific substructure (i.e. having a “*continuous AT*” loop). The second allowed for a single mismatch/unpaired base pair in the AT loop (i.e. having a “*discontinuous AT*” loop). Throughout this manuscript, these two sets of calculated probability values are referred to as “*continuous AT*” (**Table S8)** and “*discontinuous AT*” probabilities (**Tables S5** and **S7**).

Both calculations were carried out to ensure that the criteria used to obtain the probabilities of the “intermediates” allow sufficient sampling of the population, and that minute deviations from the proposed intermediates would not lead to significant changes in the probability calculations. At all transcription lengths in the studied range, the total probability difference between the *discontinuous AT* and *continuous AT* set of calculations ranged from ~ 0.0 – 0.14 (i.e. ~ 0 – 14%) before binding and 0.0 – 0.03 (i.e. 0 – 3%) after binding. Thus, the percent of RNA population available for binding is sensitive to the criteria for binding-competence, and the *discontinuous AT* criteria, which is the criteria used in calculating the presented results, sample the RNA population more efficiently than the *continuous AT* criteria.

## 4 DISCUSSION

### 4.1 The development of a ligand-induced conformational shift simulation method

RNA secondary structure prediction/folding is now a well-established field, but much less is understood about how ligands that specifically bind to RNA can induce structural changes in the population. We chose to address this problem using the well-studied SAM-I riboswitch as a model system. Many questions have been raised about the SAM-I riboswitch and riboswitches in general, (10,30,42,43,53–55) such as: How does SAM influence the base pairing competition between the P1 helix and the AT loop? What is the effect of the ligand on the RNA conformational population distribution? While many studies, including X-ray and NMR structural studies, have addressed riboswitch recognition, we aimed to understand how ligand binding perturbs the folding of the RNA, and, in turn, the downstream gene expression. While ligand binding to a conformational intermediate has been documented for at least one other riboswitch (48,56), the *yitJ* SAM-I riboswitch is unique in that binding affinity measurements are available for a significant proportion of the likely binding-competent intermediates.

To address these questions regarding the fundamental riboswitch mechanism, we combined suboptimal structure prediction with experimental binding free energies to re-calculate structural features in the population in the presence and absence of the ligand. Hence, the effect of the ligand on RNA population was simulated. Conceptually, our approach is similar to that of linked equilibria which has been a long established framework for modeling allostery and cooperativity in ligand-protein interactions (57). The study illustrates a mechanism by which the ligand shifts the equilibrium to favour a certain family of conformations, and hence leads to a transcription stop signal. Thus folding, and as a consequence gene expression, is coupled to the presence or absence of the ligand. Methods for simulating nucleotide ligand binding to RNA motifs have been previously reported (5,36). The latter method was illustrated on the binding of short RNA using oligonucleotides as ligands (5). In the current study, however, a general method to make use of binding data of small molecules was presented.

Most importantly, this is the first study to illustrate the possible role of differential/discriminatory binding between the fully folded RNA and intermediates in shaping the population distribution. Previously, aside from the aptamer forming state, few riboswitch experimental or theoretical studies took account of the possibility of binding to alternate conformers. The complex details of binding to a particular conformer, such as tertiary interactions, are intrinsically taken into account through the binding energy, since the latter was measured using an analog RNA that is constrained to form a single secondary structure, but is free to adopt a range of tertiary structures (38). By the same principle, variants in secondary structure outside of the P1/AT competition region (i.e. helices P2-P4) are implicitly present in the binding measurements.

Additionally, the method provides a means of constructing maps of probability distributions (**Figure S15** and **Figure 5**) and corresponding free energies of macrostates (**Figure S16** and **Figure S17**) before and after binding with the ligand. Such color maps can be thought of as discrete conformational and energy landscapes. Note that, **Figures S7** and **S9-S12** provide a more comprehensive picture by incorporating variants containing AT loop base pairing in conformer populations. In contrast to computing base pairing probabilities (bpp) (37), or focusing analysis on discrete lowest free energy conformations in the absence of ligand, the current method explicitly and quantitatively delineates the impact of ligand binding at the level of individual conformers. While the bpp predicts averaged values for ready comparison with popular experimental probes of secondary structure (21) in bulk, the current calculations can be compared to single molecule measurements. The experimental validation of statistical mechanical predictions provides mechanistic insights that would be lost in a purely empirical approach.

**Figure 5.**
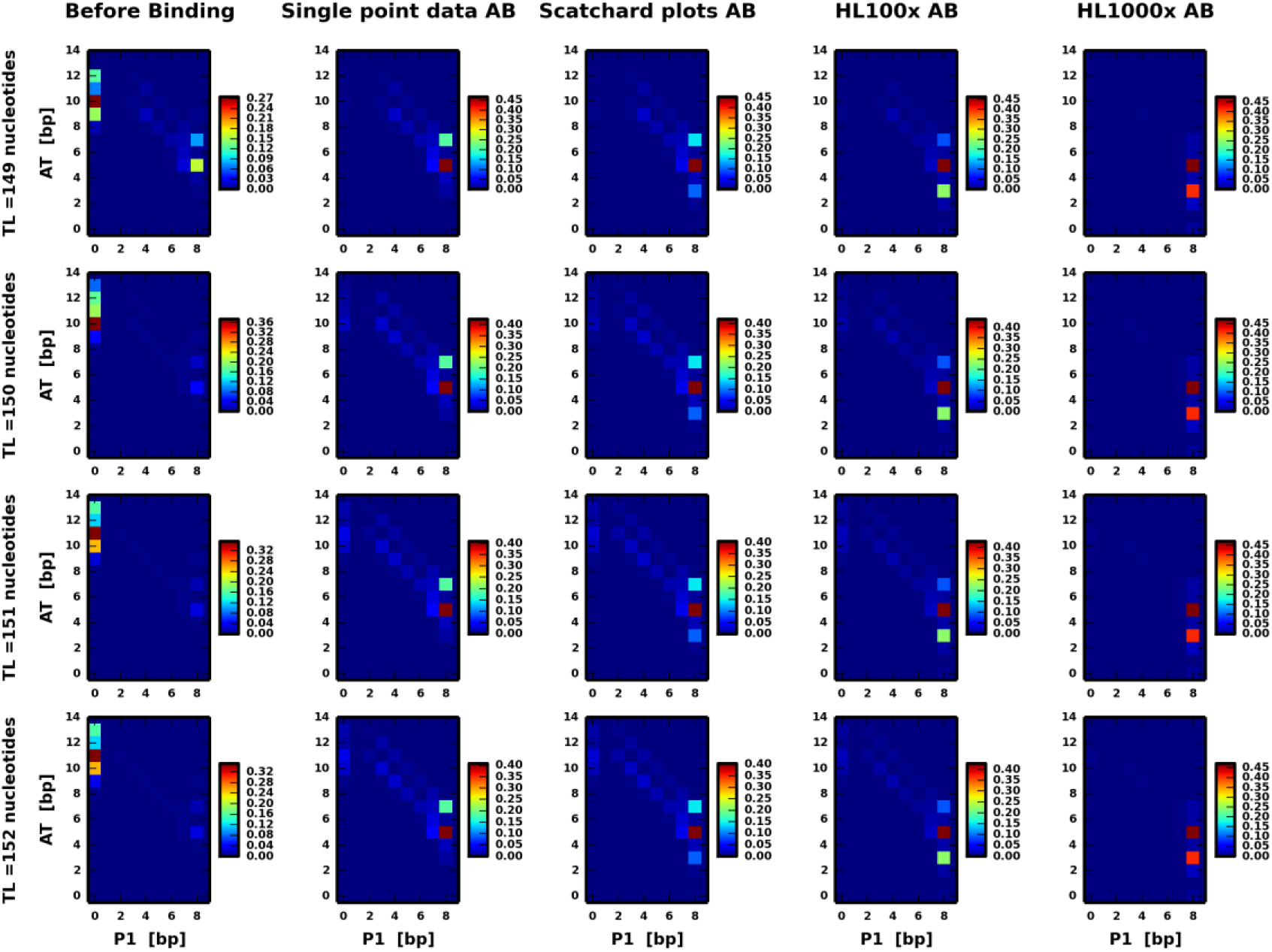
Probabilities of macrostates at transcription lengths 149-152 bases using different binding datasets. The table of color maps shows probabilities of macrostates at transcription lengths 149-152 bases (rows) and for various binding datasets used (columns). Each macrostate is represented by the sum of probabilities of conformers with a specific number of base pairs in the P1 helix and a specific number of base pairs in the AT loop (including those within the loop itself). Probabilities *before* binding are shown in the first column, and probabilities *after* binding with SAM (AB) are shown in the second - fifth for single point titration, Scatchard, HL100x and HL1000x binding datasets, respectively.

### 4.2 The differential binding between intermediates and the Aptamer is responsible for lowering the energy of the transition state

As mentioned earlier, the difference between the *P2_AT* and the *Aptamer* constructs in their binding energies is on the order of ~ −3.38 kcal/mol. The latter represents the maximum specific binding energy which is seen in the presence of a fully base paired P1 when there is no competition from the AT loop strand. The magnitude of this energy difference is approximately comparable to the favourable energy gained by the propagation of a CG pair in the neighbourhood of a GC base pair, i.e. 5’GC3’ and 3’CG5’ base pairs (−3.42 kcal/mol), at 37° C upon RNA folding (55). Additionally, to get a feeling for the magnitude of this binding energy, one may compare this value to the minimal free energy of the folding of the riboswitch at the studied lengths, which ranges from ~ −43.37 to −44.92 kcal/mol for lengths 145 and 152, respectively.

Moreover, as mentioned in the results section, the energetic discrepancy between the intermediates and the aptamer in their binding energies, as deduced from single titration point data, is on the order of ≥ 0.52 kcal/mol. The latter value is comparable to the favourable energy gained by the propagation of a GU base pair preceded by an AU base pair, i.e. 5’AG3’ and 3’UU5’ base pairs (−0.55 kcal/mol), at 37° C upon RNA folding (55). This small but significant energetic discrepancy resulting from the differential binding lowers the free energy of the *Aptamer_TS* in the presence of SAM (assuming a strand migration pathway), which likely accelerates its rate of folding (see below). The latter set of conformers possess a partially or fully unfolded AT loop.

The magnitude of the maximum specific binding energy difference obtained from the Scatchard plots was on the order of −4.76 kcal/mol, which is significantly larger than its corresponding value from single point equilibrium dialysis and comparable to the favourable energy gained by the propagation of multiple base pairs. Roughly speaking, such magnitude is comparable to ~ 1/9 of the total folding free energy of the molecule. It should be noted that binding energies from single titration point and Scatchard plots presented here were measured and calculated at room temperature, rather than 37° C. The temperature may be one of the factors contributing to the variations in values obtained by different measurements reported in the literature (38) using different techniques.

### 4.3 Reducing the energy of the aptamer transition state (*Aptamer_TS*) suggests a catalytic role of the ligand

The impact of SAM on the SAM-I riboswitch population is illustrated in the schematic diagram shown in **Figure 6**. Our results predict that even at the studied transcription lengths the unfolding of the AT loop is initiated by only a small but significant fraction (~ 2.53 - 3.63%, and may reach ~ 27.3%, as shown by the Scatchard plots binding data). A marked reduction in the probability of the *Aptamer_TS* is observed at transcription length 150 or longer, corresponding to higher energy and hence lower stability of the unfolded AT loop at these lengths. This fraction in the presence of SAM, however, is much larger than the fraction predicted in the absence of SAM (~ 0.078 - 1.49%). The latter is at least ~ 2.5 – 47 fold increase in the fraction of the population having an unfolded AT. This increase represents a reduction in the energy barrier (activation energy) required for the aptamer to cross, in order to unfold the AT loop and thereby facilitate the formation of the termination loop downstream. The reduction of the activation energy leads to the prediction that the rate of AT loop unfolding is accelerated in the presence of SAM, which can be thought of as a catalyst. Despite its significance, the percent of the functional aptamer formed is still small at this length.

**Figure 6.**
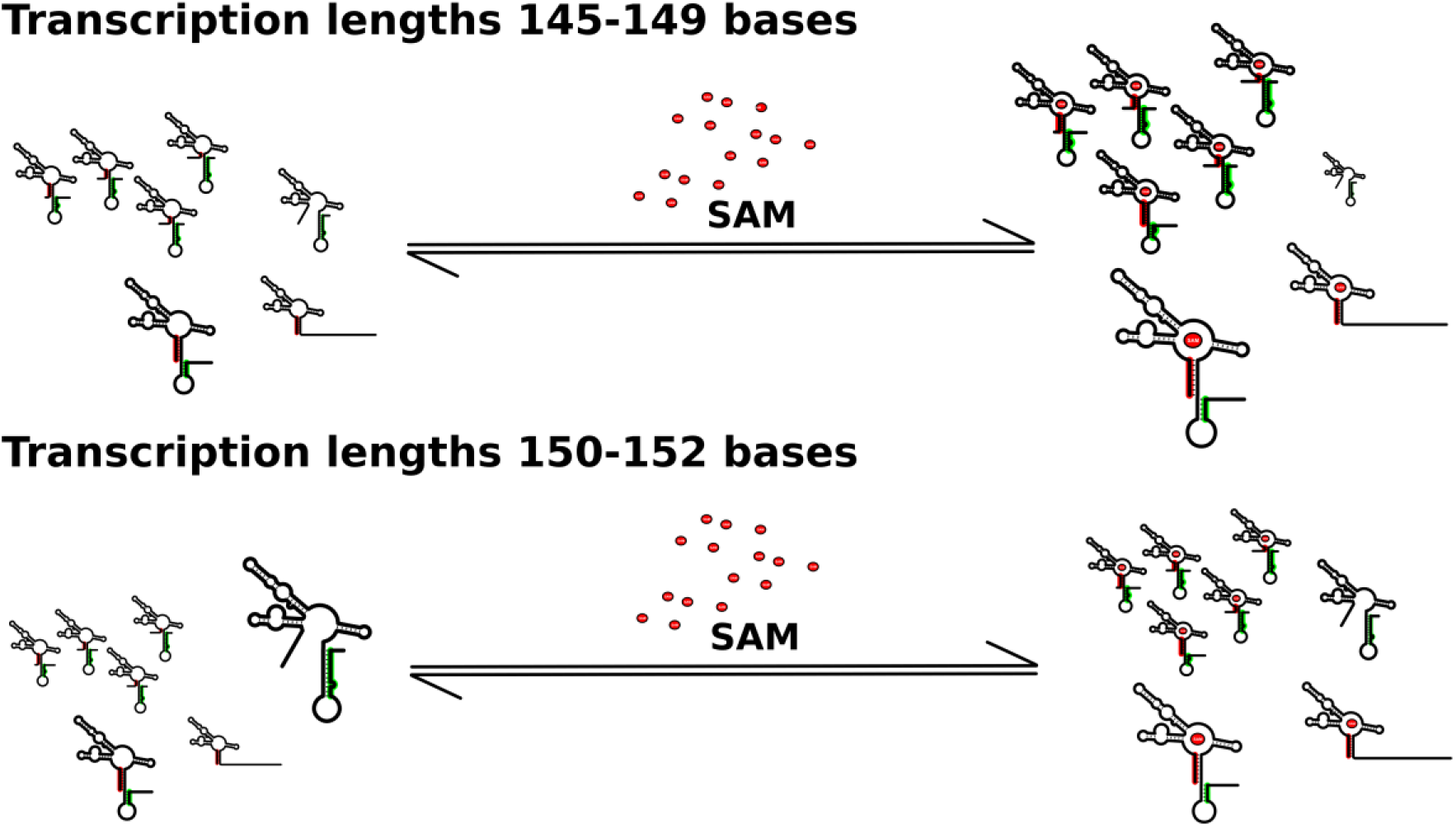
A schematic summary of the impact of SAM on the RNA population. The schematic diagrams in panels **A** and **B** illustrate the impact of the ligand (SAM) on the SAM-I riboswitch population for transcription length ranges 145-149 and 150-152, respectively. In both cases, the presence of SAM leads to conformational shifts in the population. In the absence of SAM, conformers containing the folded P1 helix either dominate or are significantly populated within the population, depending on the transcription length. Specifically, conformers containing a full length or partial P1 helix dominate the population for short transcript (panel **A**) while conformers having a base paired AT loop dominate the population for transcription length 150-152 (panel **B**). In the presence of SAM, the population shifts towards a fully folded P1 helix, which becomes dominant at all transcription lengths. Within this set of conformers containing the full P1 helix, the addition of SAM leads to an increase in the probability of the *Aptamer_TS*. The latter implies partial unfolding of the AT loop, thus paving the way for the terminator loop to be formed. In the presence of SAM, the AT loop is still significantly present in a minority population after binding. These findings suggest a strand migration mechanism. Further, these variations seen at different transcription lengths suggest that the impact of SAM is reduced at transcription length ≥ 150.

**Figure S2** illustrates examples of conformers representing the intermediates *8P1_5AT* and *4P1_9AT*, (panels **A and B**, respectively). Further, examples of binding-incompetent conformers with fully base paired AT loop are also illustrated in the same figure (**Figure S2**, panels **D - F)**. The dominance of the stable intermediate *8P1_5AT* can be understood as arising from two factors: *8P1_5AT* dominates over the *Aptamer_TS* state since the latter contains fewer total base pairs for this length of transcript. On the other hand, the preference for *8P1_5AT* over other intermediates (e.g. *6P1_7AT* or *4P1_9AT, … etc.*) that contain a similar total number of base pairs, may reflect the formation of a conformer containing an AT loop with five continuous base pairs, including a GU pair (**Figure S2A)**. Conformers with longer AT helices must necessarily form an unpaired base. Moreover, this intermediate has a slightly stronger binding free energy than other intermediates containing an equivalent number of base pairs but with only a partial P1 helix. This result is therefore expected to be sensitive to sequence variations among SAM-I riboswitches, providing a mechanism for fine tuning of riboswitch response (58).

Based on these results, we can hypothesize a transition pathway for the conversion of the predominant *8P1_5AT* conformation in the “decision window” to the terminator-containing structure. In this pathway, the *Aptamer_TS* can be considered the “transition state”. Further studies are underway to investigate how this prediction is affected when alternative parameters are invoked for treatment of dangling ends and coaxial stacking. Moreover, in the transcription complex, unlike constructs that have been used for some in vitro studies, the mRNA does not initiate at the 5’ end of the P1 helix, hence the impact of the free 5’ terminal on the stability should also be a subject of investigation.

Recently, Manz et al. (59,60) reported a study on the full-length SAM-I riboswitch using single molecule FRET. The study observed the coexistence of multiple conformers. Limited population changes were observed as a function of Mg^+2^ and SAM concentrations. In particular, conformers thought to contain the full P1 helix and the terminator helix were favoured in the presence of SAM, relative to conformers which were tentatively assigned to structures similar to those in **Figure 1**. Though these findings cannot be directly compared with our calculations for shorter transcripts, the conclusions agree with our calculations in that SAM binding increases the proportion of *Aptamer_TS* conformation, allowing the termination loop to be formed more readily than without SAM. It was also intriguing that the authors observed acceleration of folding of the SAM-bound transcription-off conformation. Although our calculations assumed thermodynamic equilibrium, accelerated formation of the same conformation is predicted if one assumes strand migration as the primary folding pathway. Thus, the hypothesized transition pathway, combined with calculations that indicate lowering of the transition state free energy barrier by ligand binding, offer a simple explanation for this experimentally observed result.

### 4.4 The role of strand migration in the absence and presence of the ligand

If SAM is absent, the P1 helix will be replaced with a fully folded AT loop, most likely through a strand migration process. In this way transcription proceeds. Previous calculations indicated that the P1 helix and terminator compete out the AT stem-loop at equilibrium for longer transcript lengths (e.g. the full length riboswitch) (37), which is not observed during transcription (45–47). It is therefore possible that the outcome is kinetically limited by RNA folding in the absence of SAM. If SAM is present, the P1 helix becomes fully base paired replacing the competing strand segment from the AT loop. The latter transition is also likely to take place through a stand migration process. Further, it was shown that adding SAM increases the (still small) probability of the unfolded AT (since we considered the *Aptamer_TS* as a mimic of this structure), paving the way for the formation of the termination loop when transcribed. Since the AT and termination loop are competing, the unfolding of the AT loop will enhance the rapid formation of the termination loop, again, through a strand migration process (10,30). Hence, strand migration is expected to play an important role in three distinct structural transitions that take place in both the absence and the presence of SAM. Since transcriptional riboswitches are thought to be kinetically controlled, strand migration is an important mechanism to accelerate folding.

### 4.5 Implications for experimental study of riboswitches and polymorphic RNA molecules

While the *yitJ* SAM-I riboswitch, to our knowledge, represents the only riboswitch system for which putative *K_d_* measurements are available for strand migration intermediates, gathering similar data for other riboswitches is not experimentally difficult. In fact, the measurements in reference (38) were performed on transcription products derived directly from PCR-amplified oligonucleotides, without any cloning. Thus, the methods here can be used to experimentally validate, in a rigorous and quantitative way, the hypothesis of ligand-induced conformational change via perturbation of the FEL for any riboswitch or other RNA systems. A handful of such studies will establish how widely the proposed mechanism can explain the function of riboswitches. Depending on the variation observed in ligand binding to intermediate conformers in riboswitch systems, it may be possible to extrapolate approximate values across, for example, SAM-I riboswitches. That would raise the possibility of predicting sequence-dependent variations across riboswitch classes, facilitating the design of a synthetic riboswitch with a specific, desired set of response characteristics.

### 4.6 Limitation of this study and future directions

Previous studies proposed the role of kinetic control for a number of transcriptional riboswitches (33,34). The proposed mechanisms suggested that the transcription rate, as well as transcriptional pausing, contribute to how the riboswitches function (61). One limitation of the method used in this study, is that kinetic control and the role of transcription rate is ignored. The assumption here is that the binding and folding rates are fast compared to transcription. With this assumption, equilibrium is reached at every nucleotide added during transcription. Local folding rates have been reported to occur on the microsecond range (although the nucleation of the P1 helix from distal strand segments may take longer) (62), whereas the polymerase transcription speed (40-90 nucleotides/s) (63) and transcriptional pausing occur on much longer time scales (seconds) (61), hence allowing the RNA-SAM complex to reach equilibrium. It has been reported that the conversion between Terminator (T) and Anti-terminator (AT) states is on the order of seconds in the presence of SAM, supporting the hypothesis that binding equilibrium may be reached (59). Moreover, calculations in this study apply only to the portion of the overall population of RNA molecules that are bound by SAM. Hence a determination of the overall conformer distribution, and therefore the proportion of terminated transcripts, must incorporate this distribution together with the distribution of unbound conformers in the correct ratios. Extensions of the method to predict titration curves are in preparation. Moreover, the statistical mechanical approach can be used within kinetic simulations, at least for specific folding pathways.

Another limitation is that the underlying folding prediction does not specifically incorporate the role of tertiary interactions such as base stacking and pseudoknot formation (43, 64–67). On the other hand, the binding free energy is based upon measurements which implicitly incorporate the range of three-dimensional structures corresponding to a given secondary structure. Thus, the energetics of the binding and the “hidden” effects on the RNA tertiary folding are treated in a “black box” as in a classical thermodynamics approach, combining them under the name “binding energy”. A fourth limitation is computational demand, particularly as the energy range for suboptimal structure calculations increases and as the length of the sequence increases. Here we have truncated the set of conformers included in the analysis such that the contribution from those that have free energies > 8 kcal/mol above the MFE are not included in the partition function. Although we have found that in this case adding contributions from additional high energy conformers does not significantly alter the result, access to high performance computing will be advantageous in wider use of the method.

The method qualitatively simulates the effect of mutants in the P1 helix and could be applied to mutants in regions that do not contact the ligand or engage in crucial tertiary interactions. In the latter cases, experimental measurement of binding to the mutant of the substructure analog, or possibly computational prediction of binding affinity, would be required.

### 4.7 Conclusions

Previously we suggested that apparently paradoxical aspects of riboswitch function, such as discrepancies between binding affinities and threshold ligand concentrations for transcription termination, could be explained by a switching mechanism involving strand migration (10,30,37). Our previous findings that SAM can bind, with reduced affinity, to proposed strand migration intermediate structures for the SAM-I riboswitch (38) raised the possibility that the ligand can accelerate switching by lowering the activation barrier. In this study, we applied a simple but rigorous statistical mechanical formalism that can form the basis for quantitative predictions and comparisons with measured populations and transition rates. For the SAM-I riboswitch, the calculations predict the observed trend of effect of the ligand on the free energy landscape, conformer population distribution and kinetics of conformational transitions. In short, they identify the stages of the transition pathway at which the ligand exerts its most important effects.

With this method, we can predict quantitatively, based on empirically based parameters for ligand binding and secondary structure free energies, how the SAM-I riboswitch ligand plays a chaperone-like role in facilitating RNA folding (10,37) and thus controlling gene expression. The strand migration process can thus be tested experimentally. The same approach is directly applicable to other riboswitches, and other RNA molecules that undergo secondary structure changes under the influence of ligand binding.

In conclusion, the simulation method used in the study yields testable predictions and, combined with hypothetical folding pathways, can be used to yield insights into kinetics. Describing ligand-induced folding of riboswitches and other RNAs using this method opens new strategies for drug design, and for the design of engineered riboswitches (30).

## 5 SUPPLEMENTARY DATA

Supplementary Data are available online.

## 6 ACKNOWLEDGEMENT

We thank Seif-Edeen Fateen (Zewail City), David Mathews (University of Rochester) and Wei Huang (Case Western Reserve University) for helpful discussions. This work was supported by Zewail City startup funds provided to the Center for X-Ray Determination of the Structure of Matter. The Center is also supported by a grant (JESOR-2015-Cycle5: Proposal ID 1008) from the Egyptian Academy of Scientific Research and Technology (ASRT).

